# Panel Perils: How Size, Fluorochrome Choices, and Unmixing Algorithms Shape Your Analysis Adventure!

**DOI:** 10.1101/2025.06.11.659156

**Authors:** Debajit Bhowmick, Timothy P. Bushnell

## Abstract

The advent of full spectral flow cytometry has enabled the development of complex panels with over 35 colors, with the latest panels reaching 50 colors (1). This capability is made possible by cytometers equipped with numerous detectors beyond those in traditional cytometers and an expanded range of fluorochromes with emission peaks across the visible spectrum. However, our observations reveal significant challenges in the current unmixing, spread prediction, and panel design methodologies. Existing tools and guidelines, largely optimized for panels with up to 20+ colors, are limited in their ability to navigate this new ultra-high-color landscape. Without improvements in unmixing algorithms, predictive tools for spread, and design strategies, researchers risk creating suboptimal panels and obtaining inaccurate results. This article aims to highlight a range of emerging challenges associated with ultra-high parameter flow cytometry, particularly for practitioners accustomed to conventional panel design and analysis. As the field advances toward increasingly complex multiparameter experiments, novel issues have surfaced—many of which were previously unrecognized. Although this work does not provide comprehensive solutions to all of these observations, it underscores the need for continued methodological development. We anticipate that ongoing research by experts in the field will yield robust frameworks to address these challenges and advance best practices in high-dimensional cytometric analysis.

**Brief description:**

1. Different unmixing/compensation algorithms can result in different biological interpretations from the same raw dataset.
2. A method to identify the optimal unmixing algorithm for accurate analysis is discussed.
3. Both panel size and specific fluorochrome combinations significantly impact population spread.
4. SSM (Spillover Spreading Matrix) values are influenced by panel size and fluorochrome combinations, which, if not carefully evaluated, may lead to misleading conclusions during panel design.

## Observations and concerns

### Concerns with Algorithms

We used the OMIP-95 (a 40 fluorochrome panel) dataset (2) to examine the impact of various unmixing algorithms on data interpretation. Specifically, we generated the unmixing/compensation matrix using single-stained and corresponding unstained samples using FlowJo (v10.10.0) Conventional, FlowJo Spectral (with the spectral box checked), FlowJo AutoSpill (with the spectral box checked), CyTEK SpectroFlo (v3.3.0), FlowLogic (solution 1.0), FCS Express (spectral, v7.24.0024), and Floreada. These matrices were applied to the same fully stained spleen FCS file (Fig. 1; see the Methods section for details). It’s worth clarifying that AutoSpill is not an unmixing algorithm (3); it is a tool for calculating single-stain spectra, used as an alternative to classic gating workflows. In FlowJo and SpectroFlo, the actual spectral unmixing algorithm is based on ordinary least squares (OLS), regardless of traditional gating or the use of AutoSpill. Primary detectors for each fluorochrome were assigned based on established criteria from the literature.

**Figure 1:**
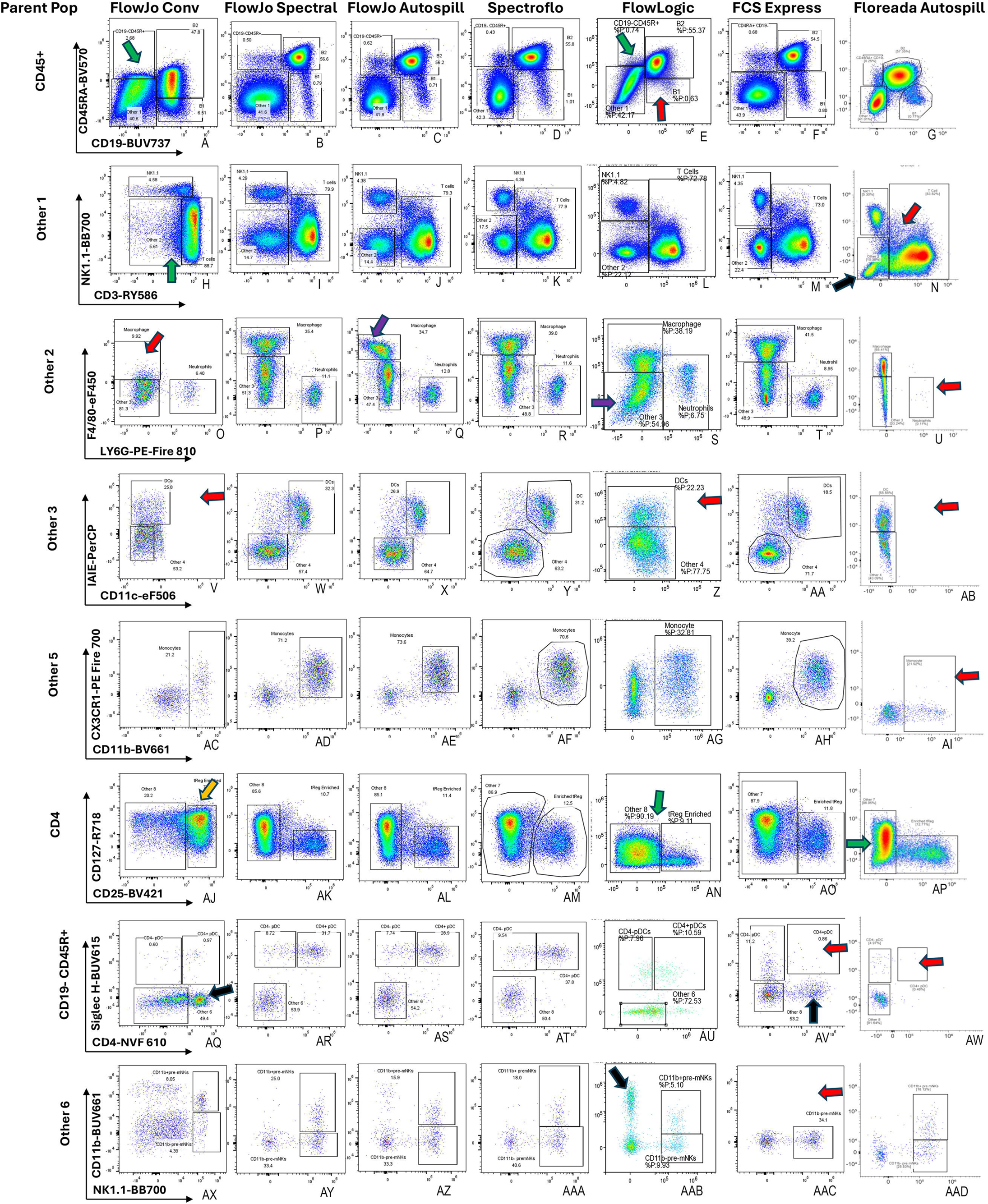
Comparison of unmixing and compensation algorithms. A fully stained spleen sample (OMIP-95) was unmixed using the same set of single-stain and unstained FCS files, processed with various commercially available algorithms (indicated at the top of each column). Analysis followed the exact gating strategy and population definitions described in published OMIP-95 (2). Parent populations are labeled on the left of each row. Colored arrows highlight specific effects observed across algorithms: Red – loss of population; Black – gain of population; Yellow – substantial change in population frequency; Green – loss of resolution; Purple – under- or over-compensated populations. All comparisons are made relative to SpectroFlo as the reference standard.

We followed the exact gating scheme described in OMIP-95 for our analysis in Figure 1. Our analysis focused on differences in unmixing outcomes relative to SpectroFlo outputs as described in OMIP-95. These differences were often substantial enough to result in inaccurate biological interpretations or misinterpretation of the data. Using 33 out of 39 fluorochromes in the panel, we identified approximately 20 instances where unmixing inconsistencies occurred. We classified these issues into five categories: (1) population loss, (2) appearance of new populations, (3) frequency changes, (4) resolution loss, and (5) incorrect unmixing (either under- or over-rectification).

For example, we observed that in cases of new populations (Fig 1, Plot AQ, AV And AAB, black arrows), inclusion of the incorrect cells in the parent gate was often the culprit, likely due to poor resolution/separation of the cells in the parent plot (Fig 1, Plot A and E, green arrow). As evident in plots B,C,D, F and G the correct cells are present in the upper left hand quadrant of each plot, enabling accurate proper analysis in row 7. When a new or unexpected population appeared, back-gating helped trace this issue; however, when populations were missed entirely (e.g., NKT cells in the Fig 1, plot N, red arrow; macrophages, neutrophils in the Fig 1, Plot O and U, red arrow; cDCs in the plot V, Z and AB, red arrow; or monocytes in the Plot AI, red arrow, or pre-mNKs in the Plot AAC, red arrow or CD4-pDC in the Plot AV, AW, red arrow), it was not possible to know with certainty if a population was present on a given plot. Notably, FlowJo AutoSpill correctly identified the NKT and neutrophil populations, while Floreada’s AutoSpill did not, even though both used the same algorithm. The observed discrepancy suggests underlying differences in algorithm implementation. This highlights the critical importance of performing rigorous quality control assessments prior to deploying specific unmixing methods.

We also identified differences in cell distribution within the double-negative populations (3rd row), with notable variation between SpectroFlo, FlowLogic, and FCS Express. Additionally, monocytes consistently appeared as double positive for CX3CR1 and CD11b in most plots (Fig 1, plot AD, AE, AF or AH), but appeared negative for CX3CR1 in Floreada’s output (plot AI) and partially on AG—a discrepancy that is particularly concerning for unknown samples or exploratory analyses. In another instance involving enriched Treg cells, Flowjo Conventional exhibited an exaggerated increase in percentage (Plot AJ, yellow arrow), whereas CD127 displayed distribution artifacts in both FlowLogic and Floreada (Plot AN and AP, Green arrow). FlowLogic claimed to achieve accuracy comparable to SpectroFlo using a mix of cell and bead-based single stains, additional methodological details might be available on their website.

To explore whether these algorithmic discrepancies were unique to OMIP-95, we conducted the same tests on OMIP-51 (4), 58 (5), and 84 (6) (containing 28-30 color panels, with the first two being conventional and the latter a full spectral panel). Interestingly, we observed only a few minor issues in the first two OMIPs and slight resolution discrepancies in OMIP-84. FlowLogic performed best across these three datasets (data not shown), suggesting that the algorithmic challenges identified may be especially pronounced in ultra-high-dimensional datasets over 30 colors like OMIP-95.

### Predicting best algorithms for unmixing

We utilized the Median Mismatch Index (MMI) matrix (7) with single-stained cells to evaluate the quality of spectral unmixing. Specifically, we applied the spillover matrix to single-stain data and quantified the median mismatch to assess algorithmic performance prior to selecting specific antibodies. For this evaluation, we employed various single stains from OMIP-095. As shown in Supplementary Fig. 1, MMI demonstrated high-quality unmixing using SpectroFlo, FlowJo Spectral, and FCS Express outputs, with few major mismatches highlighted in red. This outcome was anticipated, as the full-stain datasets performed well with these software tools.

While FlowJo AutoSpill performed well with the full stain, the corresponding MMI matrix revealed several mismatches, indicating potential issues with unmixing. Despite similar t-SNE overlays between spectral and AutoSpill outputs, subtle differences in cell distribution frequencies were observed (data not shown). Examination of the spillover matrices showed that SpectroFlo consistently produced a maximum spillover value of 1 for all fluorochromes. In contrast, AutoSpill calculations exhibited multiple spillover values exceeding 1 for several fluorochromes. When AutoSpill detects a higher spectral signature in a detector other than the primary assigned detector, it assigns a spillover coefficient greater than 1 to that channel. This results in a scaling down of the unmixed signal for the affected parameter, in contrast to gating-based "Spectral" methods, which normalize the spectral signature to the maximum-intensity detector by default. This discrepancy often occurs when the user-assigned primary detector for a fluorochrome does not have the maximum signal. For example, we identified (via Autospill) V13 and V5 as the primary detectors for four and three fluorochromes, respectively. Such assignments may explain why FlowJo Conventional performed poorly with this dataset, as conventional approaches cannot assign one detector as primary for multiple fluorochromes.

Unexpectedly, MMI indicated optimal results ("all green") for FlowJo Conventional, despite its poor performance with the full stain. Further investigation revealed that different algorithms impact the unstained populations differently, with significant variation in the relative standard deviation (rSD) of negative populations, as shown in Fig. 2 and previously described (1). Large rSD values in the negative populations skewed MMI results, yielding false-positive indications of quality unmixing. This highlights the need to carefully consider negative population spread when interpreting MMI results.

**Figure 2:**
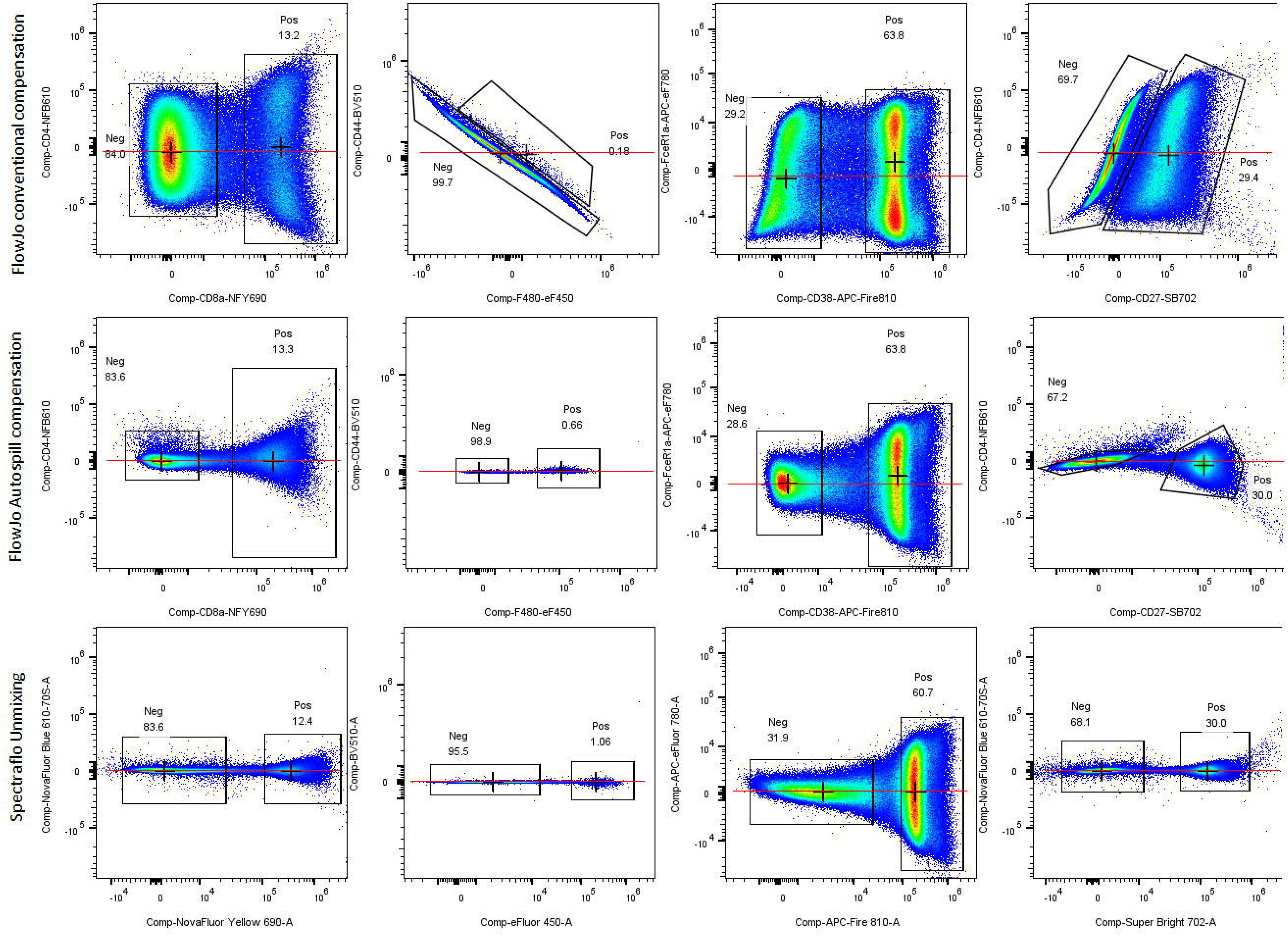
Assessment of unmixing/compensation accuracy using single-stain FCS files (OMIP-95). Single-stain FCS files were processed using 40-color matrices generated by different unmixing/compensation algorithms, indicated on the left of each row. Negative and positive cell populations were gated, and crosshairs were used to mark the MFI of each population. A red line indicates whether the MFI of the positive population (Y-axis) aligns with that of the negative population (X-axis); alignment along this line represents accurate unmixing/compensation. Deviations from the line indicate discrepancies in signal resolution.

To further explore the impact of unstained population variability, we analyzed the rSD of pure unstained cells across compensation/unmixing matrices of varying complexity. Using FlowJo Conventional, FlowJo AutoSpill, and SpectroFlo, we generated compensation matrices for 4-, 10-, 20-, 30-, 35-, and 40-color panels based on OMIP-95 single-stain data. These matrices were then applied to unstained cells. he rSD values were collected from the APC and PE-Cy5 detectors and the comparative plots are shown in Supplementary Fig. 2. Beyond the 30-color matrix, substantial distortion of the unstained population was observed—most notably with FlowJo Conventional and, to a lesser extent, FlowJo AutoSpill. In contrast, SpectroFlo preserved a well-defined unstained population. These findings indicate that as panel complexity increases, larger compensation matrices may exacerbate negative population spread, thereby hindering the resolution of dim populations in high-parameter cytometry. This distortion is consistent with previous reports on the impact of panel complexity on population spread (8). Variation in residual standard deviation (rSD) among negative populations can obscure the resolution of dim populations in large panels, even when fluorochrome combinations perform adequately in lower-parameter settings. Currently, no predictive tool exists for estimating rSD variability in negative populations, as this variability is influenced by fluorochrome selection, panel size, and cytometer hardware configuration. In a recent pre-print (9) team of Dr. Peter Mage has described the increase of the rSD in ultra large panels as Unmixing Dependent Spreading (UDS). Notably, even in 4-color panels, rSD values derived from conventional methods were higher than those observed with SpectroFlo, indicating that SpectroFlo may more effectively resolve spectral differences than conventional compensation approaches..

### Change in the rSD

As illustrated by the rising phase of the hockey-stick curve (first row, Fig. 3), this increase is strongly influenced by the specific fluorochromes included in the panel. Notably, beyond the 30-color threshold, the curve exhibits a pronounced upward inflection, with the rate of increase dependent on the particular fluorochromes added. Additionally, the number of detectors available on the cytometer contributes to this effect (data are not shown).

**Figure 3:**
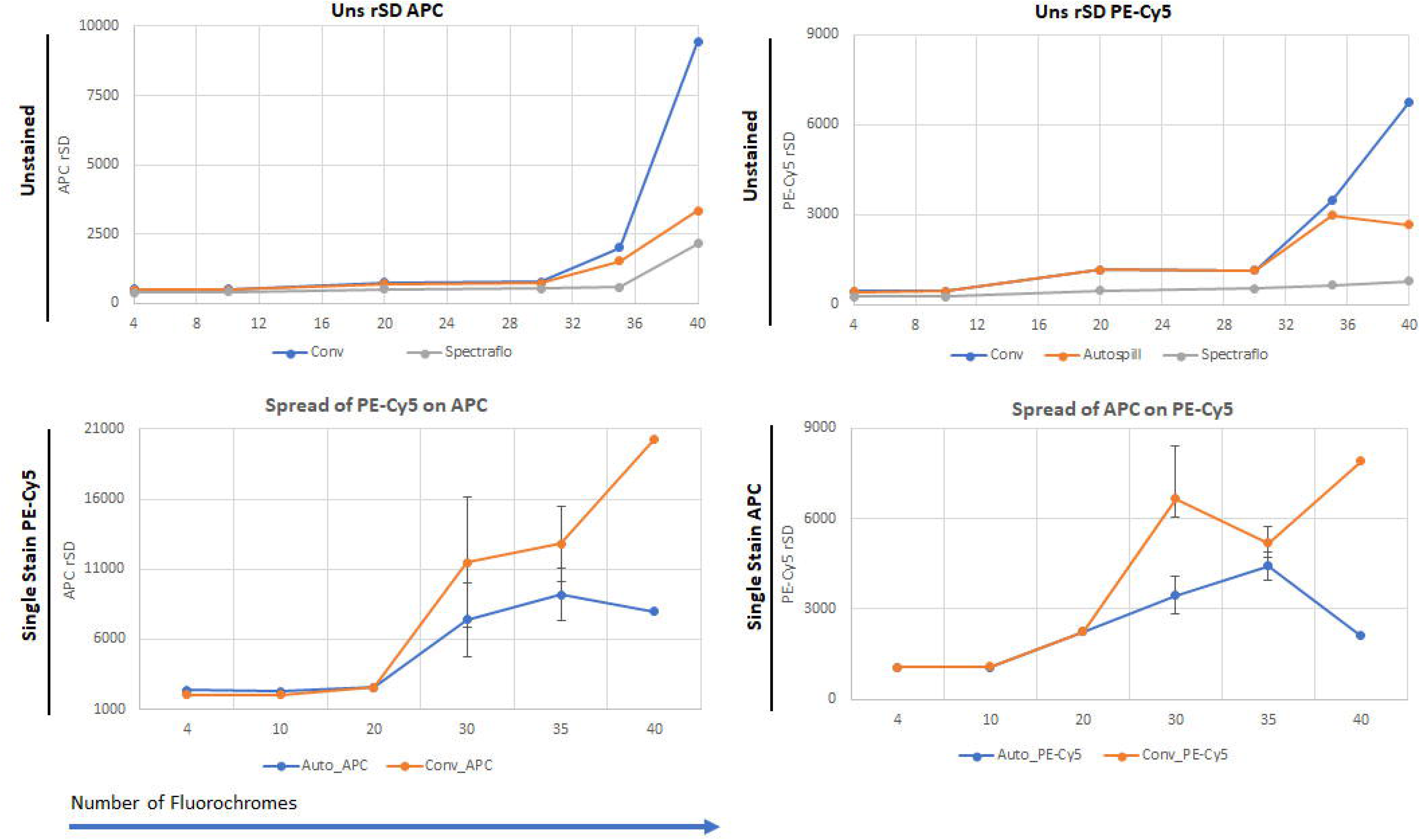
Impact of unmixing/compensation algorithms on data spread. The first row illustrates the spread of unstained cell populations, in the APC and PE-Cy5 detectors, as a function of matrix size, using FlowJo Conventional, FlowJo AutoSpill, and SpectroFlo. The second row depicts the spread of single-stained positive populations: PE-Cy5 on the APC detector and APC on the PE-Cy5 detector. Six different fluorochrome combinations were evaluated for 30- and 35-color unmixing matrices. Error bars represent variation in rSD, reflecting differences in population spread under each algorithm and panel configuration.

In the same figure, we analyzed changes in relative standard deviation (rSD) within positive population. Specifically, we evaluated single-stained APC (measuring the rSD of PE-Cy5) and single-stained PE-Cy5 (measuring the rSD of APC). The resulting plots revealed a consistent trend of increasing rSD with larger matrix sizes, independent of the unmixing algorithm used. Importantly, rSD values varied substantially based on the specific fluorochrome combinations tested; six different combinations were evaluated for the 30– and 35-color panels. These findings underscore that rSD is strongly influenced by both the size of the spillover matrix and the choice of fluorochromes.

To confirm that these findings are not unique to the OMIP-95 dataset, we analyzed an independent 50-color Aurora dataset (10). Five different fluorochrome combinations were evaluated across matrix sizes ranging from 20 to 45 colors, using three different algorithms. These matrices were applied to either unstained cell populations or single stains of APC and PE-Cy5 (rSD values shown in Supplementary Fig. 3). Interestingly, the rSD values for unstained populations segregated into two distinct groups, without exhibiting a continuous or gradual transition. This categorization appears to be primarily influenced by fluorochrome selection rather than matrix size, though the effect becomes more apparent beyond 30 parameters. In the third row of Figure 3, under SpectroFlo unmixing, the 45-fluorochrome combination highlighted in orange demonstrates an intriguing pattern: while the rSD values from the APC detector remain relatively low in both stained and unstained conditions, the same combination produces elevated rSD values in the PE-Cy5 detector. In contrast, the combination depicted in light blue exhibits consistently low rSD values across both detectors and conditions. These observations highlight the need for computational tools capable of identifying optimal fluorochrome combinations that minimize spread across all detectors. Further studies are necessary to elucidate the underlying mechanisms driving these differences and to understand their broader implications for high-parameter panel design.

A similar trend was observed in single-stained populations. Notably, the variation in rSD values for single-stained PE-Cy5 across different matrix sizes prompted further investigation into how these fluctuations might affect spread calculations using the Spillover Spreading Matrix (SSM) (11).

### Effect on Panel Design

Calculation of spread via spillover spread matrix (SSM) is a critical tool for modern panel design, whether implemented manually or using app-based solutions. To assess how matrix size influences spread, we calculated SSM values for several fluorochrome combinations from the 50-color dataset using FlowJo Spectral AutoSpill and FlowJo Conventional algorithms (Table 1 and Supplementary Table 1). As a reference point, we performed compensation using only two fluorochromes and calculated the corresponding SSM values—designated as the "ground truth" (second column, Table 1). This ground truth reflects the minimal and most accurate spread achievable when calculated from isolated single-stained controls.

**Table 1:** SSM values calculated using FlowJo Conventional. Each row represents a specific fluorochrome combination, while each column corresponds to a particular matrix combination. Compensation was performed using the FlowJo Conventional algorithm. ISS values closely matching the ground truth are highlighted in red.

|  | 2C | 50<br>C | 45_<br>1 | 40_<br>1 | 35_<br>1 | 30_<br>1 | 25_<br>1 | 20_<br>1 | 45_<br>2 | 40_<br>2 | 35_<br>2 | 30_<br>2 | 25_<br>2 | 20_<br>2 | 45_<br>3 | 40_<br>3 | 35_<br>3 | 30_<br>3 | 25_<br>3 | 20_<br>3 | 45_<br>4 | 40_<br>4 | 35_<br>4 | 30_<br>4 | 25_<br>4 | 20_<br>4 | 45_<br>5 | 40_<br>5 | 35_<br>5 | 30_<br>5 | 25_<br>5 | 20_<br>5 |
| --- | --- | --- | --- | --- | --- | --- | --- | --- | --- | --- | --- | --- | --- | --- | --- | --- | --- | --- | --- | --- | --- | --- | --- | --- | --- | --- | --- | --- | --- | --- | --- | --- |
| BB660 to PE-Fire640 | 2 | 22 | 22 | 22 | 24 | - | - | - | 22 | 23 | 23 | - | - | - | 21 | 23 | - | - | - | - | 23 | - | - | - | - | - | - | - | - | - | - | - |
| BB660 to PE-CY5 | 1 | 29 | 30 | 30 | 33 | 11 | 10 | 2 | 31 | 32 | 33 | 11 | 10 | 8 | 30 | 32 | 11 | 10 | 1 | 1 | 31 | 11 | 10 | 11 | 11 | - | 10 | 10 | 11 | - | - | - |
| PERCP to V13 | 2 | 22 | 23 | 22 | 23 | 9 | 9 | - | 23 | 22 | 22 | 9 | 9 | 8 | 22 | 21 | 10 | 9 | - | - | 22 | 10 | 9 | 9 | 5 | - | 10 | 9 | 11 | - | - | - |
| PERCP to PE-Fire640 | 2 | 53 | 52 | 52 | 54 | - | - | - | 55 | 52 | 55 | - | - | - | 53 | 53 | - | - | - | - | 53 | - | - | - | - | - | - | - | 10 | - | - | - |
| PERCP to PE-CY5 | 50 | 90 | 89 | 92 | 90 | 29 | 29 | - | 93 | 96 | 94 | 29 | 30 | 20 | 91 | 87 | 29 | 29 | - | - | 89 | 29 | 29 | 29 | 28 | - | 29 | 29 | - | - | - | - |
| BB660 to PerCP | 5 | 14 | 14 | 14 | 14 | 8 | 7 | - | 14 | 14 | 14 | 7 | 7 | 7 | 14 | 15 | 7 | 8 | - | - | 14 | 8 | 7 | 7 | 7 | - | 7 | 7 | 29 | - | - | - |
| APC-Fire750 to Alx790 | 2 | 12 | 12 | 12 | 12 | 13 | - | - | 12 | 12 | 12 | 13 | - | - | 12 | 12 | 12 | - | - | - | 12 | - | - | - | - | - | 13 | - | 8 | - | - | - |
| APC-Fire750 to APC-Fire810 | 1 | 15 | 14 | 14 | 14 | 15 | - | - | 14 | 15 | 15 | 16 | - | - | 15 | 14 | 15 | - | - | - | 15 | - | - | - | - | - | 13 | - | - | - | - | - |
| Alx700 to Alx790 | 3 | 27 | 27 | 28 | 28 | 29 | - | - | 27 | 28 | 28 | 29 | - | - | 28 | 28 | 30 | - | - | - | 28 | - | - | - | - | - | 28 | - | - | - | - | - |
| Alx700 to APC-Fire810 | 2 | 25 | 24 | 26 | 25 | 26 | - | - | 25 | 26 | 26 | 26 | - | - | 26 | 26 | 26 | - | - | - | 25 | - | - | - | - | - | 24 | - | - | - | - | - |
| ViaKrom808 to APC-Fire810 | 4 | 19 | 19 | 19 | 19 | 22 | - | - | 18 | 18 | 18 | 19 | - | - | 20 | 20 | 21 | - | - | - | 20 | - | - | - | - | - | 19 | - | - | - | - | - |
| APC-F750 to Alx790 | 20 | 27 | 27 | 28 | 28 | 30 | - | - | 27 | 28 | 28 | 30 | - | - | 28 | 28 | 30 | - | - | - | 28 | - | - | - | - | - | 28 | - | - | - | - | - |
| APC-F750 to APC-Fire810 | 7 | 39 | 40 | 39 | 37 | 41 | - | - | 39 | 39 | 39 | 37 | - | - | 39 | 39 | 41 | - | - | - | 39 | - | - | - | - | - | 35 | - | - | - | - | - |
| PE-Fire810 to BB790 | 30 | 38 | 38 | 38 | 38 | 39 | 40 | 40 | 38 | 38 | 38 | 39 | 40 | - | 38 | 38 | 38 | 40 | 41 | 41 | 38 | 38 | 40 | 40 | 40 | 31 | 38 | 40 | 40 | 31 | 31 | 31 |
| PE-Fire810 to SB780 | 1 | 17 | 17 | 17 | 17 | 17 | 17 | 17 | 17 | 17 | 17 | 17 | 16 | 1 | 17 | 17 | 17 | 17 | 17 |  | 17 | 17 | 17 | 17 | 17 | 13 | 17 | 17 | 17 | 12 | 12 | 12 |
| PE to Alx561 | 19 | 26 | 26 | 21 | 21 | - | - | - | 24 | 21 | 21 | - | - | - | 21 | 21 | - | - | - | - | 29 | - | - | - | - | - | - | - | - | - | - | - |
| PE-Fire640 to PerCP | 12 | 16 | 17 | 17 | 17 | - | - | - | 17 | 18 | 18 | - | - | - | 17 | 17 | - | - | - | - | 17 | - | - | - | - | - | - | - | - | - | - | - |
| PE-Fire640 to PE-CY5 | 5 | 38 | 39 | 40 | 40 | - | - | - | 41 | 43 | 43 | - | - | - | 41 | 40 | - | - | - | - | 39 | - | - | - | - | - | - | - | - | - | - | - |
| PE-CY5 to PE-Fire640 | 7 | 22 | 22 | 22 | 22 | - | - | - | 23 | 23 | 23 | - | - | - | 23 | 23 | 6 | 6 | 3 | 4 | 23 | - | - | - | - | - | - | - | - | - | - | - |
| PE-CY5.5 to PE-CY5 | 4 | 18 | 17 | 17 | 18 | 6 | 6 | 4 | 19 | 18 | 18 | 6 | 6 | - | 18 | 18 | - | - | - | - | 17 | 6 | 6 | 6 | 6 | 4 | 6 | 6 | 6 | 4 | 4 | 4 |
| PE-CY5.5 to NY690 | 15 | 17 | 17 | 17 | 17 | - | - | - | 18 | 19 | 19 | - | - | - | 17 | 19 | 24 | 27 | 27 | 28 | 19 | - | - | - | - | - | - | - | - | - | - | - |
| PE-CY7 to BB790 | 17 | 23 | 23 | 23 | 24 | 24 | 27 |  | 24 | 23 | 23 | 24 | 27 | - | 23 | 23 | - | - | - | - | 23 | 24 | 27 | 27 | 27 | 24 | 24 | 24 | 27 | 24 | 24 | 24 |
| BV421 to SB436 | 37 | 24 | 26 | - | - | - | - | - | 25 | - | - | - | - | - | - | - | - | - | - | - | 27 | 26 | 26 | 23 | 21 | 21 | 26 | 26 | 27 | 19 | 19 | 19 |

In most cases, we observed a substantial increase in SSM when calculated within 50-color matrices, with the extent of this increase varying by algorithm. This observation is noteworthy given that, for smaller panels, the rSD of positive population remains typically consistent pre and post compensation. This has historically supported the notion that compensation is not the source of spread, and that spread arises primarily from photon counting error at the detector level. Since the median fluorescence intensity (MFI) difference between positive and negative populations remains constant, an increase in SSM primarily reflects an increase in the rSD of the positive population. Among the tested methods, AutoSpill generally produced lower intrinsic spillover spread (ISS) values. However, we were unable to compute SSM values using SpectroFlo due to software limitations. For example, as shown in Table 1, RB755 yielded an ISS of 12 in the BV510 detector when only two fluorochromes were used, but this value escalated to 100 in the 50-color matrix—substantially impairing the resolution of double-positive populations versus RB755-positive cells. While low ISS values typically suggest that such fluorochrome pairs are suitable for resolving challenging co-expressing markers, this assumption often breaks down in larger panels, where resolution is compromised by cumulative spillover effects.

Similarly, BUV737 exhibited an ISS of 1 in the APC-Fire 810 detector at ground truth conditions. However, this value increased to 10 in some matrix combinations and reached as high as 42 in others. At present, no tool exists to reliably predict such shifts in ISS, posing a significant challenge for panel design—particularly when integrating multiple smaller panels or introducing additional fluorochromes. Without predictive insight, these modifications carry a substantial risk of reduced resolution and compromised data quality.

To further investigate, we compared 2D plots from different compensation matrices (Table 1, Supplementary Fig. 4). To further illustrate SSM discrepancies, we analyzed representative examples using both FlowJo Conventional and AutoSpill algorithms.

- **Scenario A:** The ISS of PerCP to PE-Cy5 was unexpectedly low in some panels compared to the ground truth. Specifically, in 20-, 25-, and 30-color combinations, the rSD values for both negative and positive populations were lower than the ground truth, resulting in reduced ISS values. This indicates that for certain combinations, fluorochrome pairs may achieve better resolution than in a two-color matrix. This phenomenon has not been previously documented, and no existing tools currently allow for its prediction.
- **Scenario B:** For BB755 in the BV510 detector, ISS values remained comparable to the ground truth in certain matrix configurations, as illustrated in the corresponding plots. However, for others, dramatically high ISS values were observed, which correlated with increased rSD in both positive and negative populations. Importantly, elevated rSD values in the negative population diminish the resolution of single-positive BV510 cells—an effect not captured by SSM metrics. Among the five 40-fluorochrome combinations tested, three exhibited low rSD and ISS values, while the remaining two demonstrated significantly higher spread. These findings clearly indicate that fluorochrome selection plays a more critical role in determining panel performance than the total number of fluorochromes alone.
- **Scenario C:** At ground truth conditions, the ISS of APC in the PE detector was 2; however, this value fluctuated markedly in larger panels—dropping to 0 in three matrices, rising to 10 in one, and reaching 66 in another. Typically, such low ISS values would suggest that the fluorochrome combination is well-suited for resolving low-expression or challenging antigens. However, in cases where the ISS was 0 in the 50-color and two 45-color matrices, the rSD values for both negative and positive populations were so elevated that distinction of single- or double-positive populations became effectively impossible. These findings underscore that ISS alone is not a reliable predictor of resolution, particularly in the context of high-parameter panels where spread of the negative population can obscure true signal.

These examples demonstrate that current tools, such as SSM, may be insufficient to accurately predict population resolution, particularly in high-parameter panels. Assumptions based solely on low ISS values, such as the expectation that fluorochrome pairs with ISS scores of 0 or 10 will provide adequate resolution for challenging targets, can be misleading. In practice, such pairs may perform poorly when incorporated into larger panels, leading to inadequate separation of single- and double-positive populations despite seemingly favorable ISS metrics.

To address these limitations, more advanced prediction tools are required. Spread Quality Index (SQI) (12) offers a more accurate measure of spread and, with appropriate modifications, could be adapted to predict the resolution of single-positive populations. We recommend validating the unmixing/compensation matrix using methods like MMI which offer improved insight into spread behavior. A effective spread prediction tool should include two key components: one to assess single-positive resolution and another to evaluate double-positive resolution.

Fluorochrome interactions within spectral space also play a critical role in determining unmixing performance and population resolution. For instance, although APC and Alexa Fluor 647 share similar emission peaks, they can often be successfully unmixed due to distinguishing secondary emission features excited by different lasers. However, the inclusion of fluorochromes like BV650, whose primary emission overlaps with the secondary peaks of APC and Alexa Fluor 647, can compromise unmixing performance. This interaction elevates rSD values and diminishes resolution, as demonstrated in Supplementary Fig. 4C. Such cases highlight the limitations of relying solely on SSM for panel design and underscore the need for more holistic, spectrum-aware approaches.

In conclusion, our findings emphasize the critical importance of isolating accurate positive signals for reliable unmixing—an essential step for ensuring data integrity in spectral cytometry. While panels of up to 30 colors can generally be managed effectively using most commercially available software packages, caution is warranted as panel complexity increases. For panels exceeding 30 colors, users must exercise greater scrutiny in both data analysis and biological interpretation. In such cases, cross-validating results with multiple unmixing or compensation platforms is advisable—particularly when novel or unexpected populations are identified. This multi-layered approach will help ensure that findings are robust, reproducible, and biologically meaningful in high-parameter flow cytometry experiments.

## Take home message

1. Isolating the correct positive signal is enormously important for accurate unmixing.
2. Up to 30 color panels it is relatively safe to use any commercially available software package.
3. Over 30 colors, users need to be extra careful about the biological interpretation of the data, maybe they need to use multiple packages to verify their findings, especially if they have something novel.
4. SSM does not provide reliable spread prediction for ultra high panels

## Materials and methods

### Data set

We used the following datasets FR-FCM-Z48W, FR-FCM-Z774, FR-FCM-Z6TG, FR-FCM-ZYN4, FR-FCM-ZYN4 and FR-FCM-Z4UR.

### Single Stain Processing

For all datasets, cell-based single stains were used. For OMIP-95 dataset Initial unmixing was performed in SpectroFlo using raw, unmodified single-stain FCS files. However, this approach yielded inaccurate results, necessitating preprocessing of the single-stain files prior to unmixing. As shown in Supplementary Fig. 5, we used SpectroFlo to isolate cells exclusively stained with the fluorochrome of interest, effectively excluding contributions from autofluorescence signature. This preprocessing step was conducted within SpectroFlo, as raw data exported from FlowJo were found to be scrambled. Specifically, signal data from one detector (e.g., V11) were misassigned to another (e.g., B9), with similar mis mappings occurring across multiple detectors—including shifts to UV14 and others—thereby affecting the integrity of the entire dataset.

Raw data were first plotted against time using a representative detector from each laser to identify and exclude anomalies. Manual gating was applied to remove any outliers, although in the tested datasets such events were negligible. To ensure accurate identification of stained populations, single-stain samples were analyzed in comparison to autofluorescence profiles. The detector best representing the fluorochrome signal (e.g., V12 in this case) was plotted against one or more autofluorescence-associated detectors. True positive populations appeared as distinct clusters clearly separated from the autofluorescent background. From these, the brightest stained cells (∼2,000 events per fluorochrome) were gated and exported as new, cleaned FCS files. These curated single-stain files were then used to replace the original data for all subsequent unmixing steps.

Additionally, any autofluorescence signatures were isolated in a similar manner and incorporated as virtual fluorochromes during the unmixing process. This strategy is well-described in Oliver Burton’s blog, which offers a detailed discussion of the methodology (13).

Preprocessing the single-stain files provided two key advantages. First, it significantly reduced unmixing time: FlowJo AutoSpill required approximately 3.5 hours to complete unmixing using unprocessed data, whereas the use of cleaned single-stain files reduced this time to just 6 minutes. Second, it markedly improved unmixing accuracy. Results obtained from both SpectroFlo and FlowJo closely matched the expected profiles from the OMIP-95 dataset, demonstrating high consistency and reliability.

### Comparing Algorithms

Pure and bright signals for each fluorochrome, along with autofluorescence and unstained controls, were exported in FCS format for lymphocytes, monocytes, and dead cells. These FCS files were exclusively used for unmixing and compensation comparisons. Careful matching was performed between unstained controls and their corresponding fluorochromes.

A universal negative approach was employed, using three separate unstained cell populations rather than relying on in-tube "negative" controls. These “negative” populations invariably exhibit some degree of nonspecific antibody binding, thereby violating the second rule of compensation (14F). Fluorochrome signals were identified for unmixing by collecting all positive events from the corresponding histograms.

The resultant spillover matrix was subsequently applied to the full-stain spleen FCS file for downstream analysis.

## Data availability statement

The data that support the findings of this study are available at https://flowrepository.org.

These data were derived from the following resources available in the public domain:

https://flowrepository.org/id/FR-FCM-Z48W

https://flowrepository.org/id/FR-FCM-Z774

https://flowrepository.org/id/FR-FCM-Z6TG

https://flowrepository.org/id/FR-FCM-ZYN4

https://flowrepository.org/id/FR-FCM-Z4UR

## Acknowledgement

DB is grateful to Dr. Sumanta Basu, Dr. David Novo, Dr. Bartek Rajwa and Dr. Peter Mage for their suggestion and review of the initial work. DB would like to thank Dr. Aris Kare for providing all the RAW data for OMIP-095 panel. DB likes to thank Dr. Michelle Ratliff and Dr. Mark Mannie for checking the Fig 1, DB also like to thanks Tylor for helping with analysis on FCSExpress.

## Author contribution

DB came up with the original idea, data collection, and calculation. DB and TB performed the data interpretation. DB and TB wrote the article, TB edited the final text.

---

**Supplementary Figure 1:**
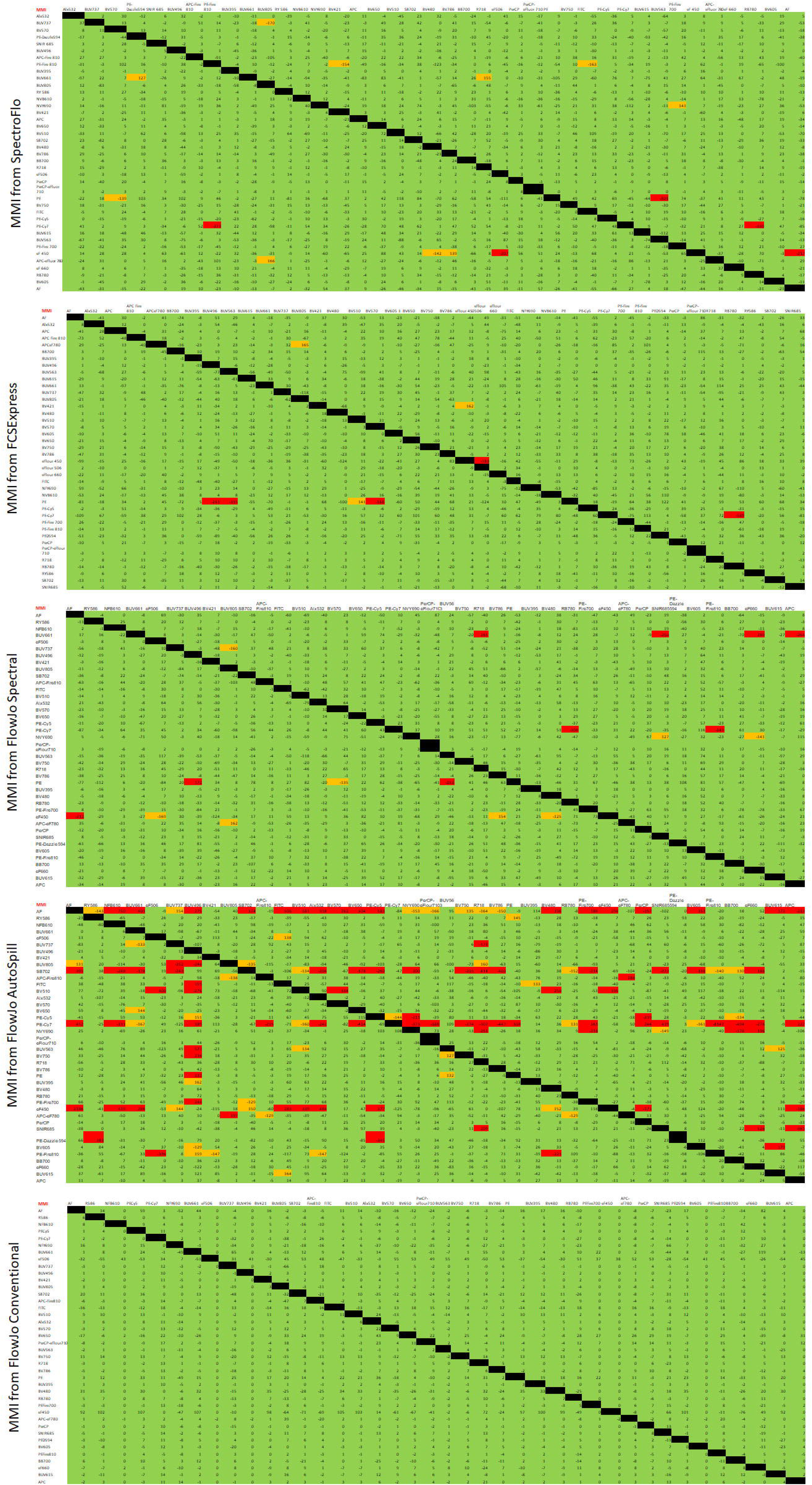
MMI analysis of 40-fluorochrome panel (OMIP-95) across different unmixing/compensation algorithms. MMI values were calculated for all 40 fluorochromes using various algorithms (listed on the left). Color-coded categories indicate the degree of mismatch between single-stained and unstained populations. Red indicates severe mismatches, where single-stained populations erroneously appear as double positives. Orange denotes moderate mismatches that may appear slightly double positive—manageable through manual gating but potentially problematic for unsupervised or automated analyses, leading to false-positive interpretations. Green represents optimal alignment, where single-stain populations match the unstained baseline. In an ideal compensation scenario, all entries in the matrix would appear green.

**Supplementary Figure 2:**
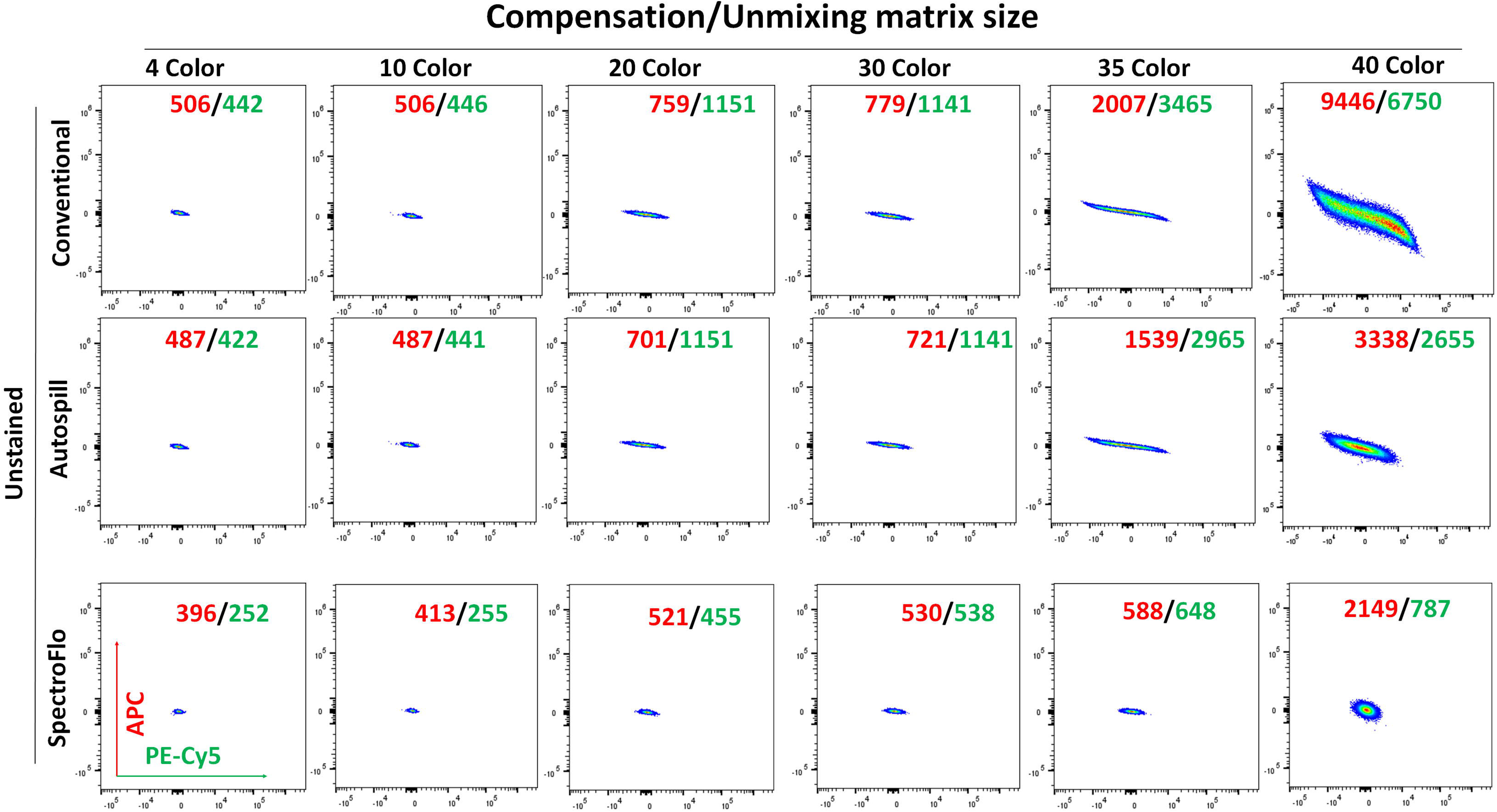
Effect of unmixing/compensation algorithms on unstained cells. Unstained cells from the OMIP-95 dataset were processed using various unmixing/compensation algorithms (listed on the left) across multiple matrix sizes (indicated at the top). Spread was quantified using rSD. In each plot, rSD values for the APC and PE-Cy5 detectors are highlighted in red and green, respectively, illustrating how detector-specific spread varies with algorithm and matrix size.

**Supplementary Figure 3:**
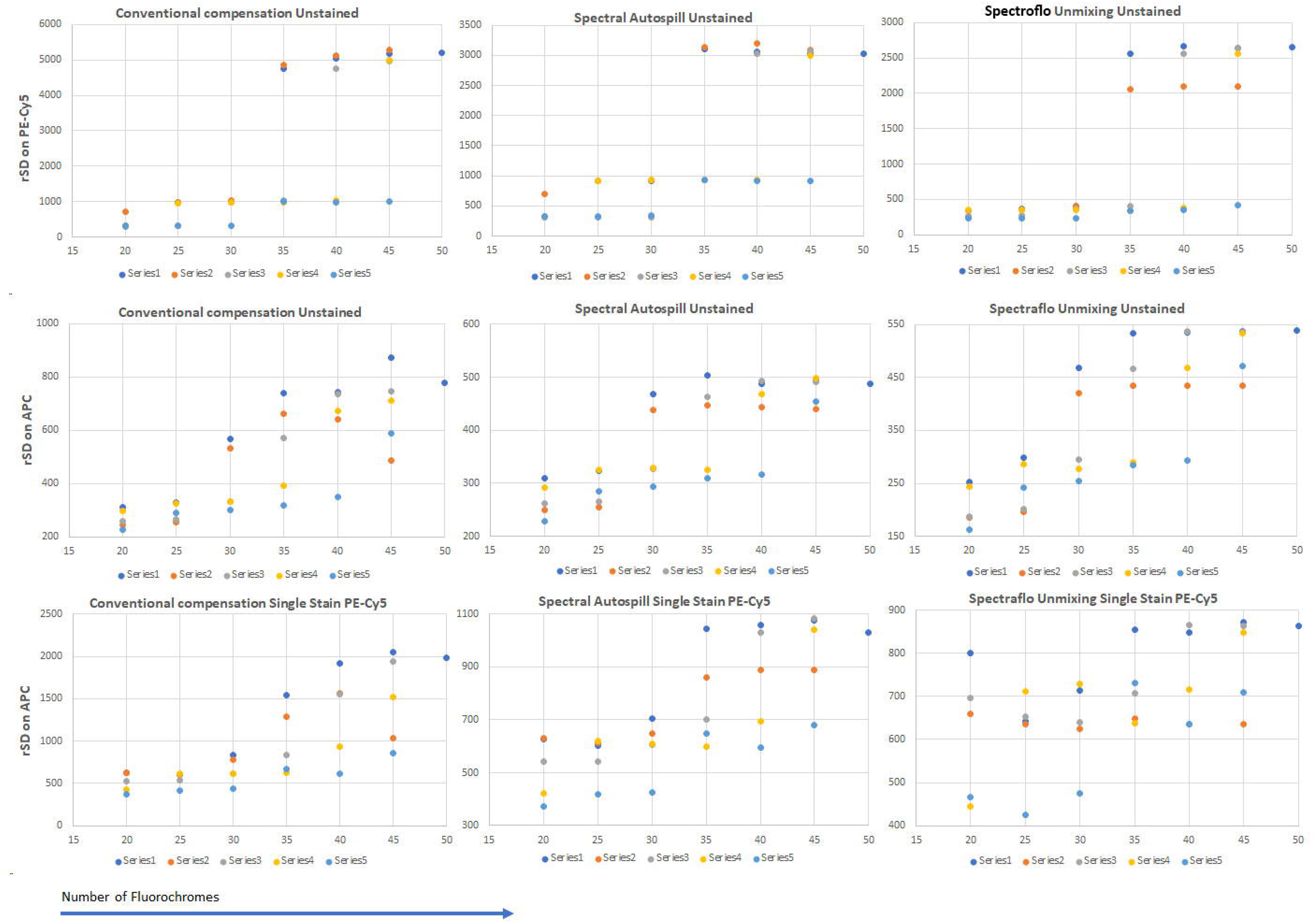
Effect of matrix size and fluorochrome combination on population spread. The 50-color Aurora dataset (1), was used to explore the impact of both matrix size and flurochome selection on data spread. The first and second rows display rSD values for unstained cells measured in the PE-Cy5 and APC detectors, respectively. Five distinct fluorochrome combinations were evaluated for a 20-color matrix, with additional fluorochromes incrementally added to generate 25-, 30-, 35-, 40-, and 45-color matrices. Each fluorochrome series is represented by a unique color; all matrix configurations are unique. The third row shows rSD values for single-stained PE-Cy5–positive cells measured on the APC detector under the same matrix conditions. This row highlights that not only matrix size but also the specific fluorochromes selected significantly influence the spread of individual fluorochromes.

**Supplementary Figure 4:**
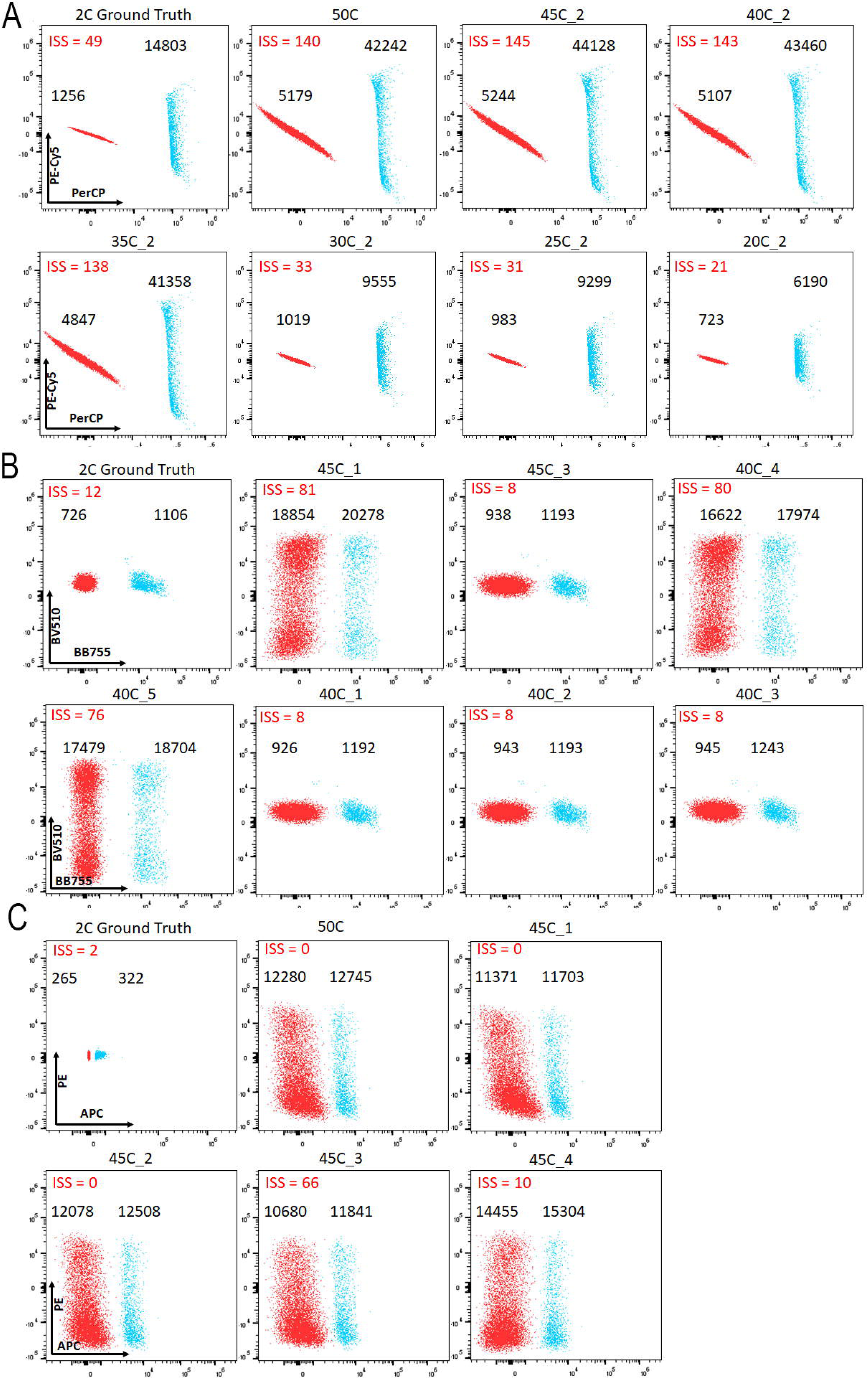
Impact of unmixing/compensation on the reliability of SSM in large panels. Unmixing/compensation influences the predictive reliability of the Spillover Spreading Matrix (SSM) in high-parameter panels. Intrinsic Spillover Spread (ISS) values are shown for three different fluorochrome combinations across multiple matrix configurations. Each plot overlays post-compensated single-stained populations with post-compensated unstained populations, using FlowJo Conventional. The rSD values are labeled in black, while ISS values are shown in red, highlighting discrepancies between predicted and observed spread.

**Supplementary Figure 5:**
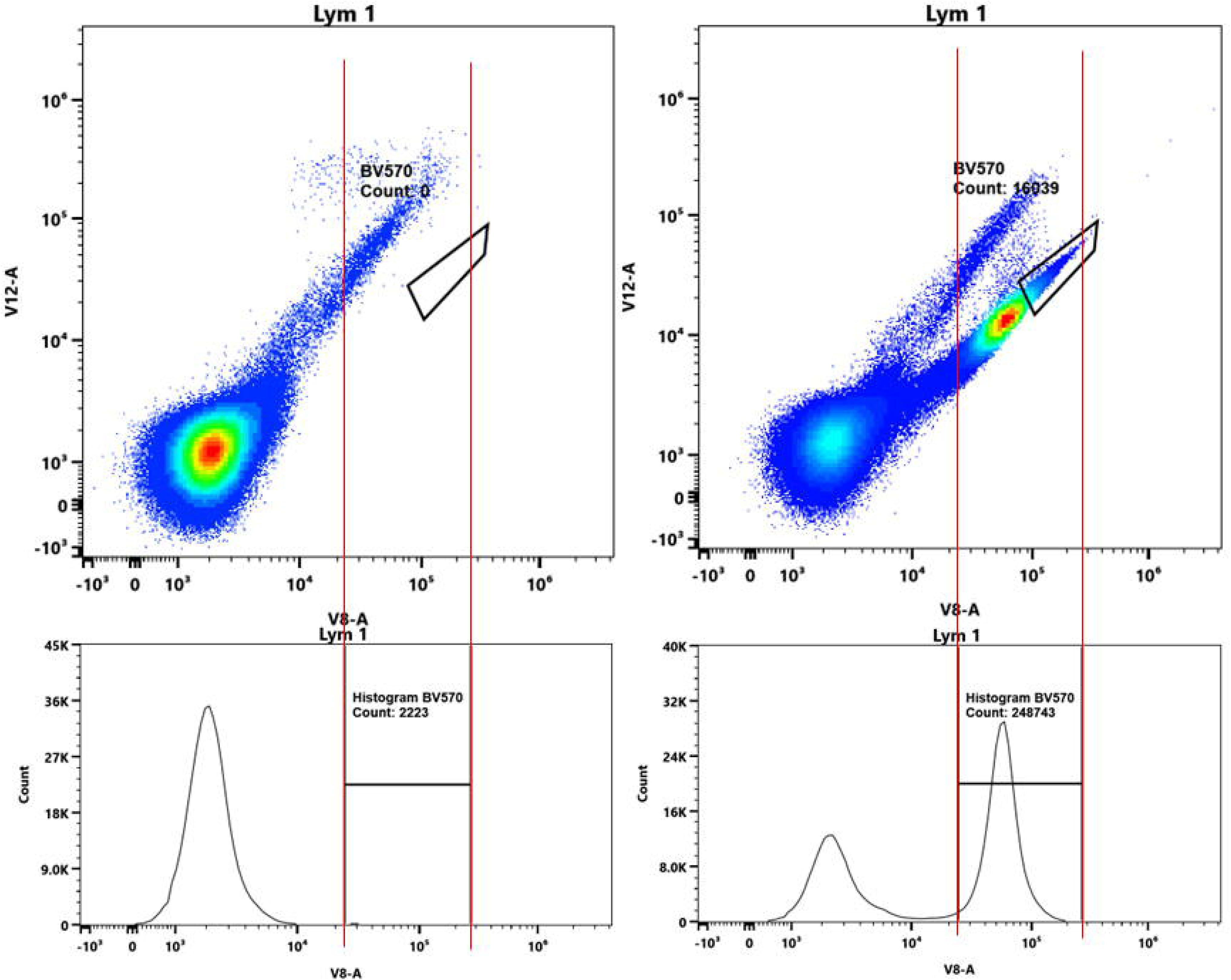
Cleaning single stains for accurate unmixing. The first row displays a dot plot distinguishing autofluorescence (AF) signals from true BV570-positive signals. Notably, highly autofluorescent cells fall within the BV570 histogram gate, despite lacking true fluorochrome staining. The second row presents a histogram combining both AF and BV570 signals, resulting in an inaccurate representation of the fluorochrome’s spectral signature. Relying solely on histogram-based gating in this context introduces multiple spectral profiles under the BV570 label, violating fundamental principles of proper compensation and unmixing (13).

**Supplementary Table 1:** SSM values calculated using FlowJo AutoSpill. Each row represents a specific fluorochrome combination, and each column corresponds to a distinct matrix combination. Compensation was performed using the FlowJo AutoSpill algorithm. That closely match the ground truth are highlighted in red, indicating minimal spread and higher compensation accuracy

|  | 2C | 50C | 45_1 | 40_1 | 35_1 | 30_1 | 25_1 | 20_1 | 45_2 | 40_2 | 35_2 | 30_2 | 25_2 | 20_2 | 45_3 | 40_3 | 35_3 | 30_3 | 25_3 | 20_3 | 45_4 | 40_4 | 35_4 | 30_4 | 25_4 | 20_4 | 45_5 | 40_5 | 35_5 | 30_5 | 25_5 | 20_5 |  |
| --- | --- | --- | --- | --- | --- | --- | --- | --- | --- | --- | --- | --- | --- | --- | --- | --- | --- | --- | --- | --- | --- | --- | --- | --- | --- | --- | --- | --- | --- | --- | --- | --- | --- |
| PE-Fire810 to SB780 | 1 | 20 | 19 | 19 | 19 | 19 | 19 | 18 | 19 | 19 | 19 | 19 | 19 | 2 | 19 | 19 | 19 | 19 | 19 | - | 20 | 20 | 19 | 19 | 19 | 14 | 19 | 19 | 19 | 14 | 14 | 15 |  |
| PE-Fire810 to BB790 | 28 | 35 | 36 | 37 | 37 | 37 | 38 | 37 | 0 | 36 | 36 | 37 | 38 | - | 37 | 37 | 37 | 38 | 38 | 39 | 35 | 36 | 38 | 38 | 38 | 29 | 36 | 38 | 38 | 29 | 29 | 29 |  |
| PE-CY7 to BB790 | 16 | 21 | 21 | 21 | 21 | 21 | 23 | - | 21 | 21 | 21 | 21 | 23 | - | 21 | 21 | 21 | 23 | 23 | 23 | 20 | 21 | 23 | 23 | 23 | 21 | 21 | 23 | 23 | 21 | 21 | 21 |  |
| Alx790 to APC-Fire810 | 20 | 34 | 34 | 36 | 36 | 35 | 22 | 21 | 34 | 37 | 36 | 36 | 22 | 21 | 35 | 36 | 35 | 22 | 22 | 21 | 33 | 33 | 22 | 22 | 22 | 22 | 34 | 22 | 22 | 22 | 22 | 22 |  |
| BB790 to SB780 | 13 | 15 | 15 | 15 | 15 | 15 | 14 | 14 | 16 | 15 | 15 | 15 | 14 | - | 15 | 15 | 15 | 14 | 14 | - | 15 | 15 | 14 | 14 | 14 | 13 | 15 | 14 | 14 | 12 | 13 | 12 |  |
| PerCP to PE-CY5 | 49 | 140 | 139 | 141 | 135 | 32 | 31 | - | 145 | 143 | 138 | 33 | 31 | 21 | 139 | 135 | 32 | 31 | - | - | 131 | 32 | 31 | 31 | 31 | - | 32 | 31 | 31 | - | - | - |  |
| APC-Fire750 to APC-Fire810 | 8 | 103 | 102 | 97 | 96 | 97 | - | - | 101 | 97 | 97 | 98 | - | - | 97 | 96 | 97 | - | - | - | 99 | 101 | - | - | - | - | 102 | - | - | - | - | - | - |
| PE-Dazzle594 to PE | 3 | 130 | 119 | 28 | 30 | - | - | - | 127 | 29 | 32 | - | - | - | 32 | 32 | - | - | - | - | 149 | - | - | - | - | - | - | - | - | - | - | - | - |
| PE-Fire640 to PE-CY5 | 5 | 75 | 75 | 79 | 74 | - | - | - | 77 | 80 | 77 | - | - | - | 78 | 74 | - | - | - | - | 69 | - | - | - | - | - | - | - | - | - | - | - | - |
| PerCP to PE-Fire640 | 3 | 96 | 95 | 98 | 94 | - | - | - | 98 | 99 | 95 | - | - | - | 96 | 94 | - | - | - | - | 91 | - | - | - | - | - | - | - | - | - | - | - | - |
| BV421 to SB436 | 50 | 80 | 82 | - | - | - | - | - | 72 | - | - | - | - | - | - | - | - | - | - | - | 83 | 69 | 69 | 45 | 41 | 42 | 70 | 68 | 47 | 48 | 46 | 46 |  |
| SBUV445 to SB436 | 3 | 39 | - | - | - | - | - | - | 43 | - | - | - | - | - | - | - | - | - | - | - | 41 | 42 | 41 | 33 | - | - | 43 | 43 | 33 | 33 | - | - |  |
| BV421 to APC-Fire810 | 1 | 42 | 44 | 42 | 42 | 43 | 10 | 10 | - | - | - | - | - | - | 42 | 42 | 42 | 10 | 10 | 9 | 41 | 42 | 10 | 10 | 10 | 10 | 43 | 10 | 10 | 10 | 10 | - |  |
| PE-CY5 to PE-Fire640 | 8 | 48 | 48 | 47 | 45 | - | - | - | 48 | 48 | 46 | 47 | - | - | 47 | 46 | - | - | - | - | 46 | - | - | - | - | - | - | - | - | - | - | - | - |
| PE-CY5.5 to PE-CY5 | 5 | 44 | 44 | 48 | 44 | 8 | 8 | 5 | 46 | 48 | 45 | 8 | 8 | - | 47 | 44 | 8 | 8 | 5 | 5 | 40 | 8 | 8 | 8 | 8 | 5 | 8 | 8 | 8 | 5 | 5 | 5 |  |
| Alx700 to Alx790 | 4 | 74 | 75 | 74 | 74 | 73 | - | - | 72 | 72 | 72 | 73 | - | - | 74 | 74 | 73 | - | - | - | 71 | 74 | - | - | - | - | 75 | - | - | - | - | - | - |
| Alx700 to APC-Fire810 | 2 | 68 | 68 | 67 | 66 | 66 | - | - | 66 | 66 | 66 | 66 | - | - | 67 | 66 | 66 | - | - | - | 66 | 67 | - | - | - | - | 68 | - | - | - | - | - | - |
| APC-Fire750 to Alx790 | 25 | 74 | 76 | 77 | 77 | 76 | - | - | 74 | 76 | 75 | 76 | - | - | 77 | 77 | 76 | - | - | - | 75 | 76 | - | - | - | - | 76 | - | - | - | - | - | - |
| NY690 to NR685 | 6 | 11 | 11 | 11 | 11 | - | - | - | 11 | 11 | 11 | - | - | - | 11 | 11 | - | - | - | - | 11 | - | - | - | - | - | - | - | - | - | - | - | - |
| ViaKrom808 to BV480 | 18 | 117 | 101 | 17 | 18 | 19 | - | - | 103 | 17 | 18 | 19 | - | - | 17 | 18 | 19 | - | - | - | 113 | 91 | - | - | - | - | 93 | - | - | - | - | - | - |
| BB755 to BV480 | 12 | 100 | 81 | 8 | 8 | 9 | 9 | 8 | 82 | 8 | 8 | 9 | 9 | - | 8 | 8 | 9 | 9 | 9 | 9 | 91 | 80 | 79 | - | - | - | 74 | 76 | - | - | - | - | - |
| BV4210 to BV480 | 3 | 175 | 156 | 3 | 3 | 4 | 3 | 3 | 155 | 3 | 4 | 4 | 3 | - | 3 | 4 | 4 | 4 | 3 | 3 | 177 | 143 | 143 | - | - | - | 151 | 146 | - | - | - | - | - |
| BV605 to BV480 | 0 | 204 | 181 | 2 | 0 | 0 | 0 | 0 | 181 | 2 | 0 | 0 | 0 | 0 | 2 | 1 | 0 | 0 | 0 | - | 215 | 163 | 164 | - | - | - | 173 | 169 | - | - | - | - | - |
| Alx561 to BV480 | 10 | 129 | 112 | 10 | 11 | - | - | - | 110 | 11 | 12 | - | - | - | 10 | 11 | - | - | - | - | 144 | - | - | - | - | - | - | - | - | - | - | - | - |
| BV605 to BV480 | 0 | 122 | 121 | 0 | - | - | - | - | 109 | - | - | - | - | - | - | - | - | - | - | - | 129 | 94 | 94 | - | - | - | 99 | 97 | - | - | - | - | - |
| PacOrange to BV480 | 2 | 116 | 114 | - | - | - | - | - | 102 | - | - | - | - | - | - | - | - | - | - | - | 122 | 86 | 86 | - | - | - | 92 | 91 | 2 | 2 | 2 | 2 |  |
| BBV48010 to BV421 | 3 | 121 | 148 | - | - | - | - | - | 111 | - | - | - | - | - | - | - | - | - | - | - | 139 | 105 | 102 | - | - | - | 116 | 103 | - | - | - | - | - |
| PacOrange to BV421 | 0 | 121 | 130 | - | - | - | - | - | 109 | - | - | - | - | - | - | - | - | - | - | - | 117 | 89 | 90 | - | - | - | 102 | 99 | 0 | 0 | 0 | 0 |  |
| BV605 to BV421 | 1 | 146 | 155 | - | - | - | - | - | 132 | - | - | - | - | - | - | - | - | - | - | - | 157 | 113 | 114 | - | - | - | 123 | 120 | - | - | - | - | - |
| ViaKrom808 to Alx790 | 8 | 97 | 100 | 102 | 101 | 105 | - | - | 97 | 104 | 104 | 104 | - | - | 101 | 103 | 105 | - | - | - | 92 | 97 | - | - | - | - | 97 | - | - | - | - | - | - |
| ViaKrom808 to APC-Fire810 | 6 | 99 | 98 | 95 | 94 | 95 | - | - | 99 | 98 | 98 | 98 | - | - | 95 | 100 | 95 | - | - | - | 93 | 99 | - | - | - | - | 100 | - | - | - | - | - | - |
| PerCP to BV4213 | 2 | 45 | 45 | 45 | 42 | 14 | 13 | - | 46 | 45 | 42 | 14 | 13 | 12 | 45 | 42 | 14 | 13 | - | - | 42 | 16 | 14 | 13 | 13 | - | 15 | 14 | 14 | 14 | - | - | - |
| APC to PE | 2 | 0 | 0 | 59 | 60 | - | - | - | 0 | 57 | 59 | - | - | - | 66 | 59 | - | - | - | - | 10 | - | - | - | - | - | - | - | - | - | - | - | - |
| PE-Fire640 to PE | 1 | 78 | 76 | 52 | 55 | - | - | - | 74 | 49 | 50 | - | - | - | 54 | 54 | - | - | - | - | 95 | - | - | - | - | - | - | - | - | - | - | - | - |
| BV421 to PE | 2 | 79 | 79 | 59 | 60 | - | - | - | - | - | - | - | - | - | 64 | 60 | - | - | - | - | 89 | - | - | - | - | - | - | - | - | - | - | - | - |

